# scDynOmics: An Optimized Transformer Model for Representation Learning from Single-Cell Multiomics

**DOI:** 10.64898/2026.02.28.708160

**Authors:** Gang Yu, Timothy J.S. Ramnarine, Johanna Klughammer, Simon W. Mages

## Abstract

As foundation models have become increasingly prevalent in several fields for multiple purposes, pretraining models with single-cell transcriptomic data has gained significant interest. Although existing single-cell foundation models have demonstrated that transformer-based designs can be applied to various biological tasks, they do not show consistently competitive performance compared to much simpler approaches while requiring much more resources in terms of data and compute. Here, we introduce scDynOmics, a pretrainable multiomics-capable transformer for representation learning from single-cell data. The model is motivated by gene regulatory networks without excluding unknown interactions between genes and adopts a Linformer-style attention mechanism to scale to coding-genome wide multimodal inputs. Pretraining on paired single-cell transcriptomic and chromatin accessibility profiles yields compact high-fidelity embeddings that represent cellular states and developmental dynamics. For versatile application, scDynOmics employs low-rank adaptation modules, enabling parameter-efficient fine-tuning for downstream tasks. We demonstrate that scDynOmics outperforms existing single-cell foundation models by a large margin and achieves or surpasses state-of-the-art performance compared to simpler approaches, while revealing interpretable factors driving developmental trajectories and perturbation responses that simpler approaches cannot provide. Overall, scDynOmics is an efficient, scalable, flexible, and interpretable framework for cellular representation learning and deciphering cellular heterogeneity and dynamics.

## 1 Introduction

Recent advances in high-throughput next-generation sequencing (NGS) technologies have transformed our ability to measure molecular profiles at scale. This enabled a revolution in the study of cellular heterogeneity by single-cell RNA sequencing (scRNA-seq), gene expression profiling at single-cell resolution, offering unprecedented insights into the diversity of cell states[1]. Current spatial transcriptomics (ST) methods build on this by preserving the spatial organization of cells, allowing researchers to analyze cellular states within the context of tissue architecture[2]. Furthermore, single-cell multiomic sequencing (scMultiomics) integrates transcriptomic and epigenomic information, such as chromatin accessibility profiles (scATAC-seq), from the same cell, enabling a more comprehensive characterization of cellular states[3]. Integrative, multimodal analyses of scMultiomics data have revealed intricate regulatory mechanisms and identified key lineage transitions during cellular development and in disease contexts[4, 5]. Analytical methods in both multiomics and mono-omics, for example, RNA velocity applied to scRNA-seq to infer the direction of cellular state change, have proven effective for dissecting complex biological processes[5, 3, 6]. Despite these advances, traditional analytical approaches fall short in comprehensively capturing the complex patterns present in the datasets influenced by technical noise[7], motivating the development of scalable and efficient tools to extract deeper biological insights from these high-dimensional data.

The advent of foundation models has sparked a revolution across the machine learning community, with pretrained Transformer[8] models consistently demonstrating their robustness and effectiveness across various domains, including natural language processing (NLP)[9][10] and computer vision (CV)[11]. Inspired by these advances, similar models have been applied to gene expression matrices derived from scRNA-seq data. Unlike autoencoders, which had early success in tasks such as denoising scRNA-seq data[12, 13], pretrained Transformers offer improved generalization by leveraging a large-scale data corpus during pretraining and fine-tuning for various downstream tasks[14]. Existing approaches, often pretrained with millions of single-cell gene expression profiles, have achieved notable success in tasks such as cell type classification, perturbation effect prediction, and gene regulatory network (GRN) inference[14]. However, many of these models are constrained by the quadratic complexity of standard Transformer architectures, which limits their ability to scale to long input sequences spanning the full gene space.

To address this challenge, many current single-cell foundation models rely on a selected subset of the most informative features to generate initial representations[15, 16]. While effective in reducing computational cost, such feature selection strategies may inadvertently exclude biologically important genes, particularly in contexts where regulatory influence is specific to a cell type or condition. A few methods attempt to achieve coding-genome scale input with optimized attention mechanisms[17, 18] or alternative architectures like state space models[19, 20]. Nevertheless, the potential of customizing attention mechanisms to advance scalability for single-cell data that explicitly reflect the structure of biological regulation remains largely unexplored.

The increasing interest in processing long sequence data, such as video data[11], has led to the development of various efficient attention mechanisms. Some of the most intuitive approaches belong to the class of sparse attention mechanisms[21, 22] which approximate the full attention matrix by selecting representative subsets of the input and processing them with standard attention mechanisms. In the context of single-cell data, where the regulatory importance of individual genes is often unknown *a priori*, such selection strategies may inadvertently exclude critical information and thus limit the model’s performance. Current methods mostly focus on the most expressed or most variable genes, and limited alternative mechanisms have been explored[14]. Another class of methods reduces the computational complexity of attention from *O*(*N*^2^) to approximately *O*(*N*) by modifying the attention computation itself, as exemplified by architectures such as Linformer[23], Performer[24], and LinearAttention[25]. Within the single-cell domain, scBERT[17] has applied the Performer architecture, while CellFM[18] utilizes retentive networks[26]. These are the few models enabling the encoders to process full coding-genome scale input. Besides the algorithmic optimizations mentioned above, several recent works have also explored hardware-level optimizations, leading towards kernel optimization methods such as FlashAttention[27, 28] and SAGEAttention[29]. Published methods, such as scGPT[16] and scPRINT[30], have already explored corresponding optimizations but still struggle to achieve coding-genome scalability.

In this project, we propose scDynOmics, a pretrainable model for scalable cellular representation learning with optimized attention mechanisms and biologically motivated design principles. Specifically, we adopt sparse attention and linear attention mechanisms to enable the model to process multimodal coding-genome scale input. Building on the concept of GRNs[5], we hypothesize that transcription factor (TF) mediated regulation induces a low-rank structure in single-cell data, consistent with the low-rank adaptability assumption in Linformer[23]. Accordingly, in each layer of scDynOmics, we project the input into a latent space representing activating regulons in cells, which typically constitute a much smaller subset of the genome, while retaining unprojected query vectors to preserve whole-genome expressivity. This design choice allows the model to capture essential regulatory relationships while maintaining computational efficiency. Moreover, to support parameter efficient fine-tuning (PEFT) for various downstream tasks (e.g., cell fate prediction, annotation), we integrate low-rank adaptation (LoRA) modules[31, 32, 33] into the pretrained model. Leveraging this fine-tuning strategy, scDynOmics requires significantly reduced computational resources as well as smaller datasets for adapting the model to specific biological contexts. Although the use of projected latent features within the scDynOmics model makes a direct interpretation of attention matrices challenging, we implement a gradient-based explanation mechanism to elucidate the model’s behaviors, enabling the extraction of biologically meaningful insights. Finally, through extensive evaluation, we demonstrate that scDynOmics not only achieves or surpasses state-of-the-art performance in transfer learning but also provides interpretable predictions that reveal complex developmental trajectories and spatial perturbation effects.

## 2 Results

### 2.1 scDynOmics

An overview of the scDynOmics framework is presented in Fig. 1. Single-cell profiles, such as gene expression matrices, are high-dimensional, typically comprising *L* ≈ 20, 000 genes in human and mouse cells[14]. Standard Transformer architectures process such inputs with a self-attention mechanism that scales quadratically with input length (*O*(*L*^2^)), rendering them computationally prohibitive for coding-genome scale applications.

**Figure 1:**
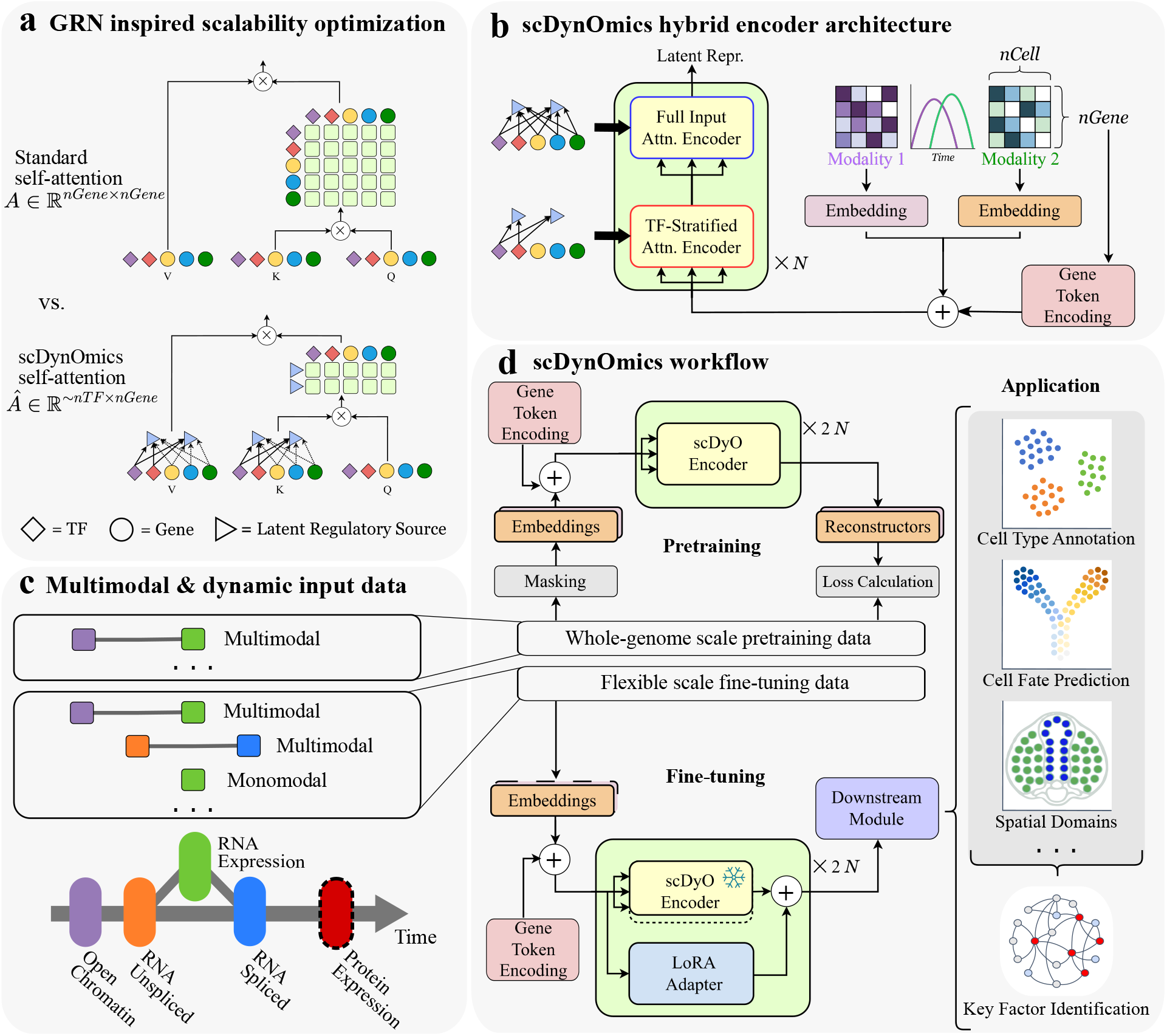
Overview of the scDynOmics framework. **a**, The gene regulatory network (GRN) inspired scalibility optimization of the attention mechanism. In standard self-attention with an input length at the full coding-genome scale *L*, the attention matrix *A* has quadratic complexity based on the number of genes. In the optimized linear attention mechanism of scDynOmics, the attention matrix is approximated via a low-rank projection to dimension *l*, corresponding to the number of transcription factors (TFs). **b**, Hybrid encoder architecture. The model stacks alternating layers of TF-stratified attention encoder (TF-Encoder), which constrain ***K*** and ***V*** projections to documented TFs, and full input attention encoder (Full-Encoder), which utilize learnable dense projections across the full feature space (Methods). Ideally, the input consists of paired modalities profiling genes’ temporal dynamics. **c**, The paired multimodal or monomodal inputs to scDynOmics. **d**, The scDynOmics workflows. The pretraining phase utilizes a masked-input reconstruction objective to learn generalizable cellular representations. The fine-tuning phase integrates adapter modules for parameter-efficient adaptation to downstream tasks. The learning objective in fine-tuning is heavily task-dependent and part of the specific downstream module.

To overcome this bottleneck, scDynOmics adopts a Linformer-style linear attention mechanism, approximating the full rank attention matrix via a low-rank projection to dimension *l* ≪ *L*, where *l* approximates the number of activating TFs (Fig. 1a, Methods). This design aligns scDynOmics’ latent attention space with the biological structure of regulatory logic. Recent studies on single-cell GRNs view cellular systems as sets of regulons centered on TFs to study biological progressions, such as differentiation[5, 34, 35, 36]. We propose that a *O*(*lL*) attention space can effectively capture key regulatory relationships governing cellular states and dynamics while ensuring computational tractability.

To optimize this projection, we developed a hybrid encoder architecture (Fig. 1b, Methods). Strictly constraining projections to known TFs ensures biological consistency but may fail to capture unannotated regulatory factors. Conversely, learning unconstrained projections allows for flexible exploration but ignores valuable biological priors, potentially requiring redundant computational efforts to rediscover known TFs. To balance these considerations, scDynOmics stacks alternating layers of *TF-Encoders*, which guide the model towards known TFs, and *Full-Encoders*, which allow exploration of the entire coding-genome.

During multimodal pretraining, scDynOmics distills biologically meaningful cellular embeddings by capturing the dynamic responses of these regulons to cell intrinsic and external signals (Fig. 1c, d). To operationalize these embeddings for diverse biological inquiries, scDynOmics integrates adapter modules that enable PEFT on downstream tasks with minimal computational overhead. Furthermore, we couple this adaptation strategy with a custom gradient-based attribution framework (Methods), allowing us to deconvolute the model’s outputs into interpretable signatures.

### 2.2 Optimization of Model Architecture and Pretraining Efficiency

To establish a robust foundation for the scDynOmics framework, we first sought to determine the optimal model architecture that balances computational efficiency with representational capacity.

Utilizing the comprehensive mouse scMultiomics pretraining corpus (*N* = 752, 155 cells; *L* = 22, 452 genes; Fig. 2a, Methods), we conducted a systematic grid search to optimize key hyperparameters: the latent dimension *l* for the linear attention mechanism, and the encoder depth and width. We first analyzed the distribution of expressed TFs across the corpus, observing that all cells expressed fewer than 700 TFs (Fig. 2b). Consequently, we capped the search space for the latent dimension *l* at 700. Models were pretrained (*n* = 3 independent runs) using the Masked Input Prediction (MIP) objective with a 10% validation hold-out (Methods). Performance was evaluated based on the cross-entropy loss of reconstructing masked values (local feature learning) and the entire input (global structure preservation). We found that a projection dimension of *l* = 500 offered an optimal trade-off between both aspects of reconstruction accuracy and computational cost (Fig. 2c). Notably, we compared our proposed hybrid encoder architecture against a model using exclusively Full-Encoders (500 FGP). Despite a marginal advantage in masked reconstruction, the 500 FGP model underperformed in reconstructing the full input and required significantly more parameters (139M) compared to the hybrid model (78M). Therefore, we selected the hybrid encoder architecture for all subsequent experiments.

**Figure 2:**
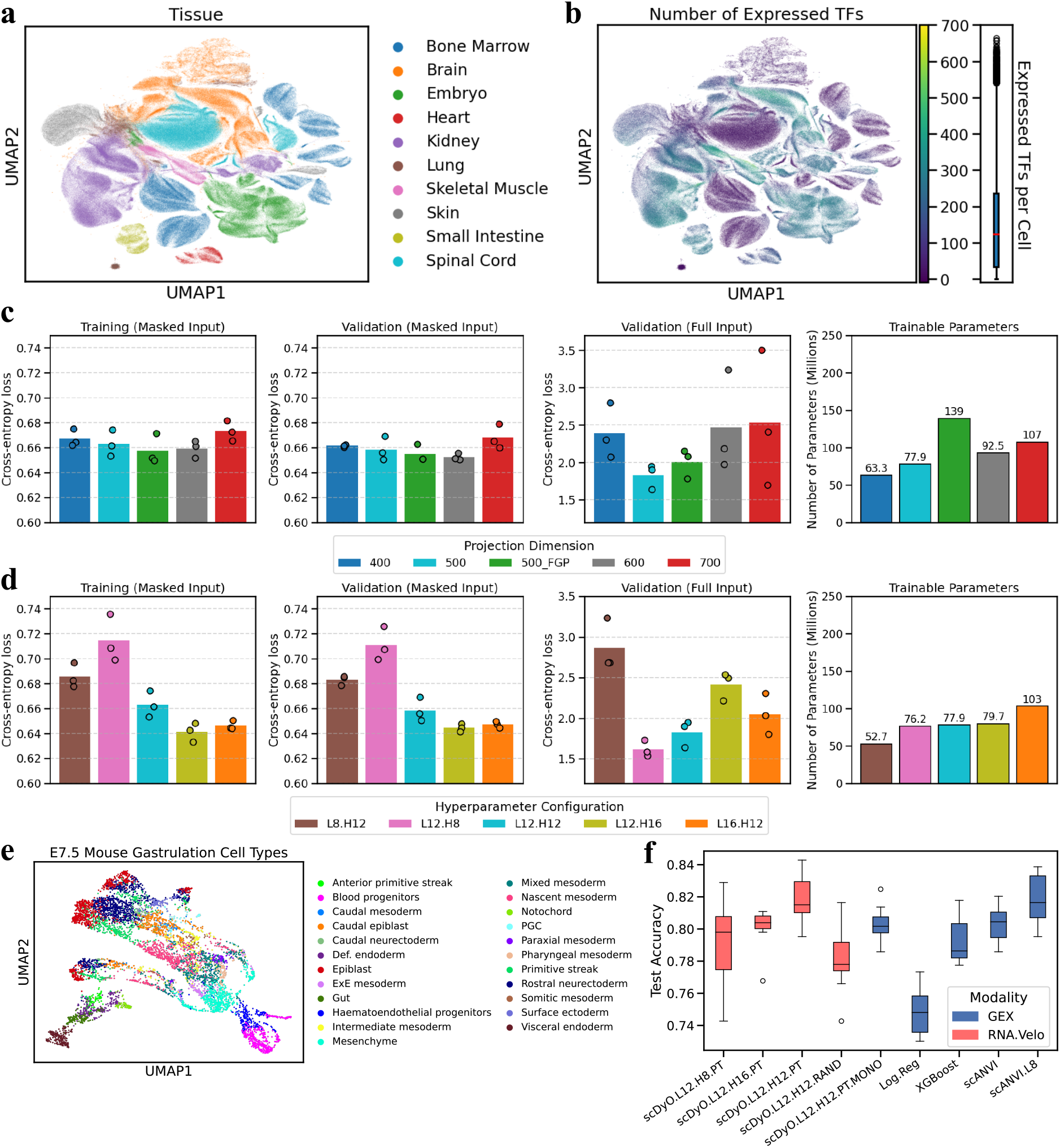
Hyperparameter optimization and pretraining validation. **a**, UMAP embedding of the transcriptomic profiles from the curated mouse scMultiomics corpus used for pretraining colored by tissue. **b**, Same UMAP embedding as in **a**, colored by the number of expressed transcription factors (TFs; left) and boxplot of the distribution of expressed TFs per cell (right) illustrating the coverage of the search space for the latent attention dimension *l*. **c**, Training loss (left) and validation reconstruction loss (for masked input indicating local feature learning, mid left, and for full input indicating global structure preservation, mid right) with parameter efficiency comparison (right) across varying latent dimensions *l* and model architectures. The full genome projection variant for latent dimension 500 (500_FGP) replaces the TF-Encoders in scDynOmics by Full-Encoders. **d**, Training loss (left) and validation reconstruction loss (for masked input indicating local feature learning, mid left, and for full input indicating global structure preservation, mid right) with parameter efficiency comparison (right) across different model hyperparameter configurations (number of layers and number of attention heads). **e**, UMAP embedding of the mouse gastrulation dataset utilized for fine-tuning the cell type classification task colored by celltype. **f**, Downstream classification accuracy on the mouse gastrulation dataset, benchmarking the selected L12H12 architecture against alternative configurations and non-pretrained baselines, fine-tuned (scDynOmics) or trained (non-pretrainable baselines) either on regular mono-modal gene expression count data (GEX) or multi-modal spliced and unspliced count data (RNA.Velo; color).

We further examined the impact of model depth and width (Fig. 2d). We identified three configurations, all utilizing 12 layers (L12) with approximately 80M parameters, that achieved comparable pretraining performance. While the 16-head configuration (L12H16) excelled in masked reconstruction, the 8-head configuration (L12H8) proved superior in full input reconstruction. The 12-head configuration (L12H12) provided a balanced performance across both metrics.

To empirically resolve this trade-off, we evaluated the transfer learning capabilities of these three configurations by transitioning from the scMultiomics pretraining approach to an alternative fine-tuning setup (Methods). Specifically, we assessed whether models pretrained on total RNA and *pan-promoter* ATAC-seq could implicitly capture RNA expression dynamics by re-aligning the model input in a multimodal RNA velocity framework during fine-tuning: we applied a temporal remapping where the *pan-promoter* ATAC-seq modality was replaced with unspliced pre-mRNA counts, and the gene expression modality was replaced with spliced mRNA counts. Because unspliced counts reflect nascent transcription and spliced counts represent mature mRNA available for translation, this remapping allows the model to capture active transcriptional dynamics in a conceptually similar manner to the scMultiomics pretraining setup, enabling pretrained representations to adapt effectively. As the experimental validation of this approach, we utilized a downstream cell type classification task on a mouse gastrulation dataset (*N* = 7, 187 cells; *L* = 22, 439 genes)[37] (Fig. 2e).

In a 10-fold cross-validation setting with test folds stratified by cell type, the L12H12 model yielded a higher median classification accuracy 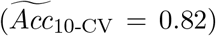 than both the L12H8 and L12H16 variants (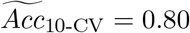 each) (Fig. 2f, models L12.H8.PT, L12.H12.PT, L12.H16.PT). Therefore, we selected the balanced L12H12 architecture as our default configuration.

To benchmark scDynOmics against established single-modality methods (Methods), we adapted the classification task to use total RNA counts rather than partitioned spliced and un-spliced transcripts. In this experiment, the multimodal L12H12 model outperformed standard linear (Logistic Regression, 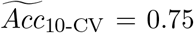) and decision tree-based baselines (XGBoost[38], 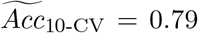). Furthermore, it had comparable performance to the representative Variational Autoencoder[39] (VAE)-based benchmark, scANVI[40] after hyperparameter optimization (median accuracy 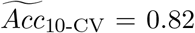), demonstrating state-of-the-art performance (Fig. 2f, models Log.Reg, XGBoost, scANVI.L8).

We next evaluated the importance of scMultiomics pretraining for this RNA velocity inspired task. Despite the change in modality, pretraining increased the median accuracy from the non-pretrained baseline 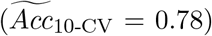 (Fig. 2f, model L12.H12.RAND). Crucially, we tested whether scDynOmics genuinely transfers dynamic biological insights across modalities, rather than trivially learning to aggregate spliced and unspliced counts. Fine-tuning the pre-trained model solely on total RNA (monomodal configuration) resulted in a diminished accuracy 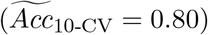 (Fig. 2f, model L12.H12.PT.MONO) compared to the multimodal setup. This performance gap confirms that scDynOmics successfully captures and transfers complex dynamical relationships between distinct multimodal paradigms.

### 2.3 Efficient Transfer Learning from Specialized Pretraining Data

To address scenarios where massive, general-purpose pretraining corpora are unavailable or limited in covering the area of interest, we investigated the efficiency of scDynOmics when pretrained solely on a modest, topic-specific dataset. We utilized the curated human scMultiomic immune cell dataset (*N* = 48, 067 cells; *L* = 20, 492 genes; Methods) to pretrain the model on a specialized immunological context. The model was subsequently fine-tuned to predict cell type annotations using the monomodal scRNA-seq peripheral blood mononuclear cell (PBMC) profiles of the widely utilized Zheng68k dataset[41], consisting of *N* = 68, 450 cells; *L* = 13, 709 genes (Fig. 3a). We evaluated multiple scDynOmics configurations using 10-fold cross-validation with test folds stratified by cell type, benchmarking them against prominent pretrained foundation models (scBERT[17], Geneformer[15], CellFM[18]) and established machine learning baselines (Logistic Regression, XGBoost, scANVI[40]). Consistent with best practices for optimizing cell type annotation[42, 43], we preprocessed the fine-tuning dataset to retain the top 2, 000 highly variable genes (HVGs) selected with Scanpy[43]. In addition to providing all features, this feature selection was applied to scDynOmics, CellFM, and the baselines. However, scBERT and Geneformer were evaluated adhering strictly to their native input requirements to ensure a fair and rigorous comparison. Specifically, scBERT was provided only with the default full coding-genome scale inputs, and Geneformer was evaluated utilizing its signature rank-value encoding of top 2000 non-zero expressed genes.

**Figure 3:**
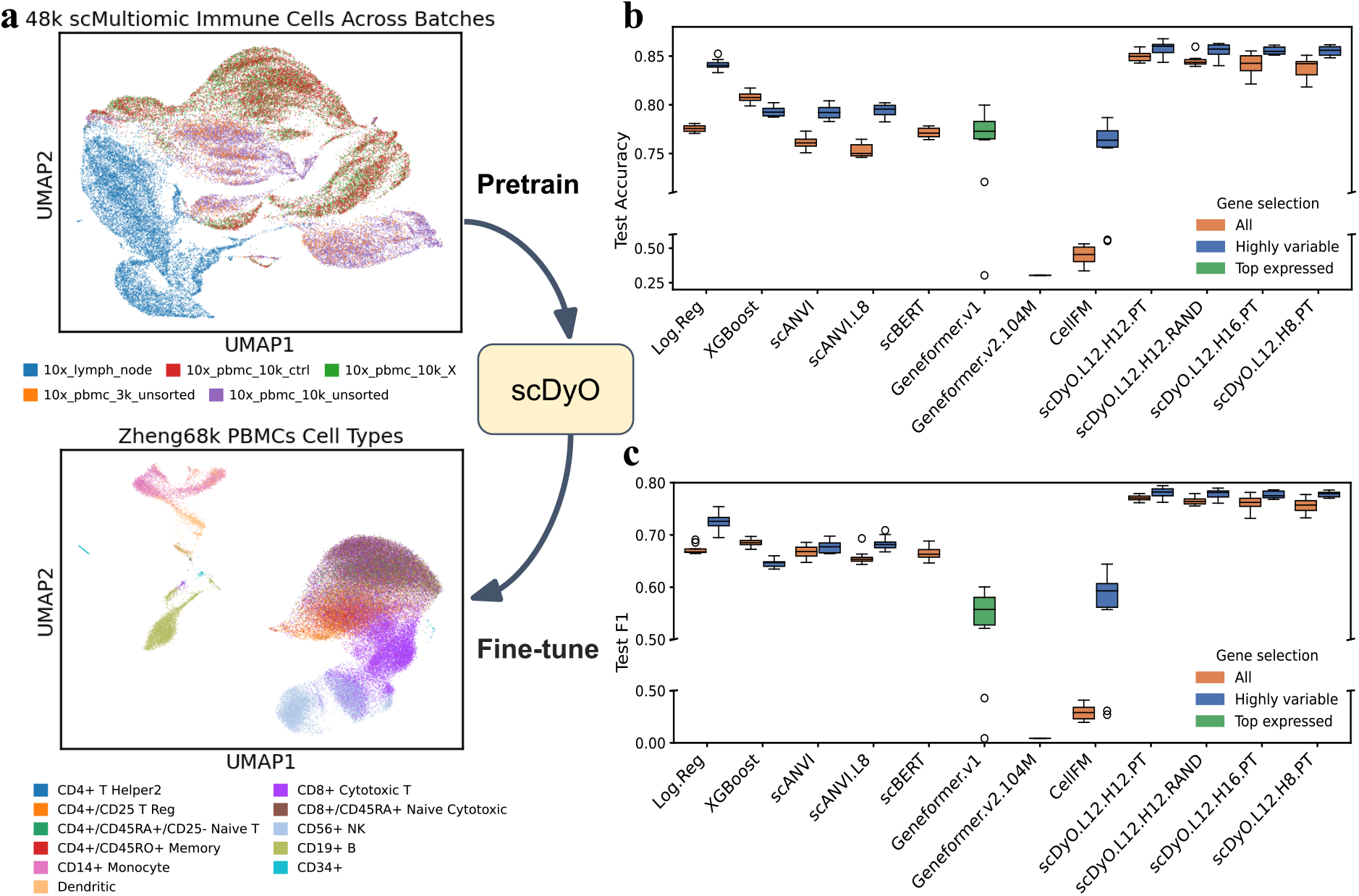
Evaluation of transfer learning efficiency on human PBMC datasets. **a**, Schematic of the experimental workflow: pretraining is conducted on a specialized, limited corpus of human scMultiomic immune cell profiles (top: UMAP embedding colored by batch), followed by fine-tuning on the monomodal Zheng68k scRNA-seq peripheral blood mononuclear cell (PBMC) dataset (bottom: UMAP embedding colored by cell type). **b**,**c**, Boxplots comparing classification accuracy (**b**) and weighted F1-scores (**c**) across 10-fold cross-validation. scDynOmics configurations reach at least comparable performance with the baseline models (Logistic Regression and XGBoost) while outperforming existing foundation models.

Despite the limited scale of the pretraining corpus, scDynOmics achieved state-of-the-art classification performance with or without the HVG selection step, surpassing pretrained foundation models and reaching performance levels comparable to or exceeding baseline methods. With the non-pretrained L12H12 model trailing only slightly behind the pretrained L12H12 counterpart, the actual benefit of pretraining on this small corpus was limited with the performance difference comparable to other effects, like choosing different hyperparameters for the pretrained scDynOmics. Notably, while a better performance on cell type classification on HVGs compared to using all features is common among the benchmarked methods, scDynOmics keeps its comparatively strong performance also without feature selection. This indicates that, beyond the benefits of transfer learning, the intrinsic architecture of scDynOmics could capture cellular semantics efficiently, suggesting its potential as a robust tool even when pretraining resources are limited.

### 2.4 Uncovering Regulatory Drivers of Developmental Transitions

Beyond accurate classification, a critical utility of pretrained single-cell models lies in their capacity to translate generalizable patterns obtained from large-scale data into specific biological insights. To validate the interpretability of scDynOmics, we applied the selected L12H12 model pretrained on our large curated mouse corpus from Section 2.2 to a held-out scMultiomics dataset characterizing the differentiation trajectory of mouse embryonic stem cells (mESCs)[44] (Fig. 4a). We specifically focused on the subtle transition window between 48 hours (h48) and 52 hours (h52) post-differentiation induction. To ensure a rigorous evaluation of the model’s fine-tuning transferability, *N* = 1, 000 cells with *L* = 18, 285 genes from each time point were randomly held out from the pretraining phase. For all benchmarked models, the resulting dataset was partitioned into fine-tuning (90%) and validation (10%) sets. After fine-tuning the pretrained model for binary discrimination between these temporal states, we computed the discriminative specificity score (**d**_*c*_; Methods) for each feature to isolate the key factors driving this transition.

**Figure 4:**
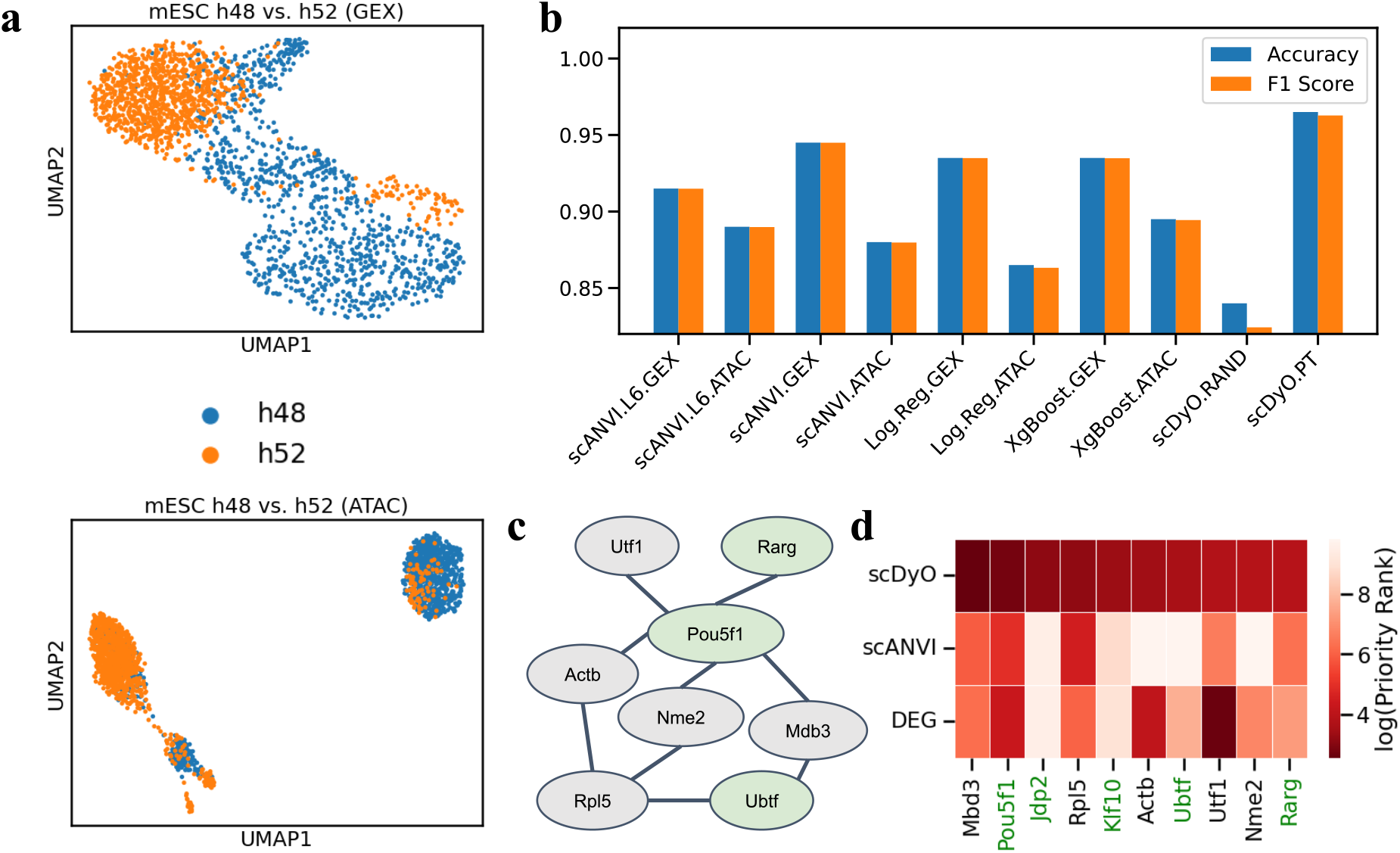
Interpretability of mESC differentiation dynamics. **a**, UMAP embedding of mouse embryonic stem cells (mESCs) at 48 (h48) and 52 hours (h52; color) post-differentiation induction, based on gene expression (top) and ATAC profiles (bottom). **b**, Benchmarking of classification performance (Accuracy and F1-score) against baseline methods on the held-out test set. **c**, Key regulon reconstructed from the top 50 discriminative features identified by scDynOmics using the STRING[45] interaction database. Documented transcription factors are highlighted in green. **d**, Heatmap comparing priority rankings (color) of features (columns) between scDynOmics attribution, differentially expressed genes (DEGs) from standard analysis, and interpretation of the scANVI model (rows). Key regulators like *Pou5f1* show high model attribution despite low priority assigned by other methods, illustrating the model’s ability to capture complex regulatory signals.

In comparison with baseline models, the pretrained scDynOmics model achieved state-of-the-art classification performance with validation accuracy and F1-score both around 0.96 (Fig. 4b). However, the model’s value extends beyond prediction metrics. Our gradient-based analysis identified a compact set of high-ranking regulatory factors, most notably *Pou5f1* (*Oct4*), *Jdp2*, and *Mbd3* (Fig. 4c-d). While *Pou5f1* is a well-established master regulator of pluripotency[46], the identification of *Mbd3* and *Jdp2* highlights the pretrained model’s sensitivity to complex regulatory dynamics: *Jdp2* has been reported in several studies as a key factor in facilitating somatic cell reprogramming[47, 48], demonstrating its critical role in modulating cellular pluripotency; *Mbd3*, a core component of the NuRD complex, plays a pivotal role in chromatin remodeling and the exit from pluripotency, yet its expression levels often exhibit only modest changes during such short temporal windows [49, 50]. Consequently, neither standard differential expression analysis nor interpretation of the scANVI model prioritized these genes (Fig. 4d).

The ability of scDynOmics to pinpoint key regulatory factors that standard differential expression analysis and baseline model interpretation fail to identify suggests that the pretrained model offers unique, biologically grounded insights into developmental transitions that are not accessible through linear statistics alone.

### 2.5 Deciphering Progenitor Fate from Mature Signatures

We next investigated whether the learned cellular representations capture the key signals governing the developmental continuum. To achieve this, we evaluated scDynOmics pretrained with the mouse scMultiomics corpus on a cell fate prediction task using a spatial transcriptomics (Slide-seq) dataset of the mouse embryo[51], retaining *L* = 18, 324 genes. We formulated a “time-reversed” generalization problem: models were trained to distinguish mature cell types, specifically those located in the differentiated neural tube (*N* = 2, 288 Slide-seq beads) and somites (*N* = 616 beads), and subsequently tested on their ability to predict the fate decisions of upstream progenitors (*N*_*presomitic*_ = 278 beads; *N*_*neuromesodermal*_ = 149 beads) located in the tailbud region (Fig. 5a). This experimental setup rigorously tests the model’s capacity to identify early lineage-priming signals that precede overt differentiation, leveraging ground-truth annotations associated with spatial coordinates.

**Figure 5:**
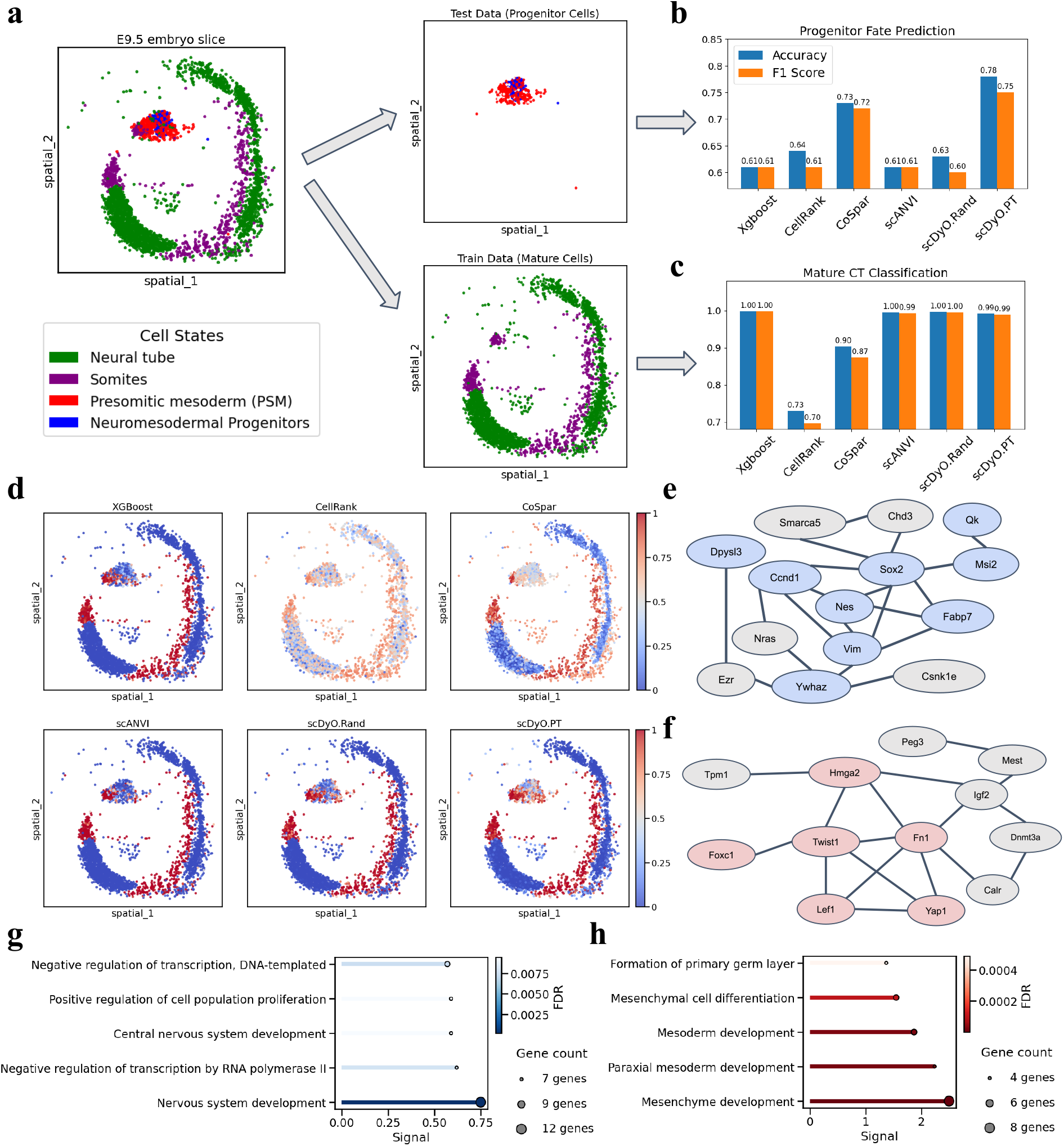
Cell fate prediction in spatial transcriptomics. **a**, Schematic of the “time-reversed” prediction task: models are trained on mature cells (Neural Tube vs. Somites) and tested on upstream progenitors in the tailbud. **b**,**c**, Classification metrics on the progenitor test set (**b**) and the mature cell training set (**c**) for all benchmarked methods. **d**, Spatial distribution of prediction confidence scores (color) across mature and progenitor populations for all benchmarked methods. **e**, Key regulon reconstructed from the top 50 discriminative features identified by pretrained scDynOmics for predicting Neuromesodermal Progenitors, utilizing the STRING[45] interaction database. Genes associated with the Gene Ontology (GO) term “nervous system development” are highlighted in blue. **f**, Key regulon reconstructed for predicting Presomitic Mesoderm. Genes associated with the GO term “mesenchyme development” are highlighted in red. **g**,**h**, Top 5 enriched GO terms (rows) of the top regulon for the neural (**g**) and somitic (**h**) lineage, showing the Benjamini-Hochberg FDR (color), the STRING Signal score (x axis), and the number of overlapping genes between the top 50 discriminative features and that GO term (symbol size).

While most benchmarked methods achieved near-perfect accuracy in classifying the mature training profiles, their performance dropped significantly when generalizing to the progenitor test set (Fig. 5b-c). scDynOmics, however, achieved the highest predictive accuracy of 0.78 with a macro F1-score of 0.75. This outperformed the second-best method, CoSpar[52], an optimal-transport (OT) based approach explicitly designed for lineage inference, which achieved an accuracy of 0.73 and a macro F1-score of 0.72. We also evaluated CellRank[53, 54], another prominent trajectory inference framework. However, it exhibited suboptimal performance on both mature and progenitor classification in this setting, potentially attributable to its distinct reliance on Markov chain modeling derived from OT kernels. Interestingly, scDynOmics exhibited reduced predictive confidence on progenitor profiles, similar to CoSpar, compared to the relatively higher confidence on mature cells (Fig. 5d). This observation might reflect the inherent biological plasticity and transitional nature of the progenitor state. Overall, these results suggest that pretrained scDynOmics offers robust predictive power regarding progenitor cell fates.

To validate the biological basis of these predictions, we analyzed the key regulons driving scDynOmics’ classification of progenitors as either neuromesodermal (Fig. 5e) or presomitic mesoderm (Fig. 5f). Gene Ontology (GO) enrichment analysis of the top 50 predictive features revealed terms highly relevant to the respective lineages. For instance, terms such as “nervous system development” were enriched for neural predictions (Fig. 5g), while “mesenchyme development” was enriched for somitic predictions (Fig. 5h). These findings further confirm that the model’s predictive capability is driven by biologically coherent developmental programs rather than confounding artifacts.

### 2.6 Resolving Spatial Heterogeneity in Perturbed Embryos

Finally, we investigated the model’s capacity to resolve complex spatial phenotypes resulting from genetic perturbations (Fig. 6a). We utilized Slide-seq profiles (*L* = 11, 300 genes) from wild-type (WT) and *Tbx6* knockout (KO) mouse embryos. In wild-type development, *Tbx6* is essential for specifying paraxial mesoderm; its absence prevents cells bilateral to the central neural tube from differentiating into somites, causing them to adopt a neural fate and form “ectopic neural tubes”[51]. Although these ectopic cells retain some transcriptomic signatures of their somitic origin, their gene expression profiles are also highly consistent with central neural tube cells[51]. The WT data consists of *N*_*somitic*_ = 103 somitic and *N*_*neural*_ = 87 neural annotated Slide-seq beads, while the KO data contains *N* = 305 “neural” beads including both regular central and ectopic neural tube cells (Fig. 6b).

**Figure 6:**
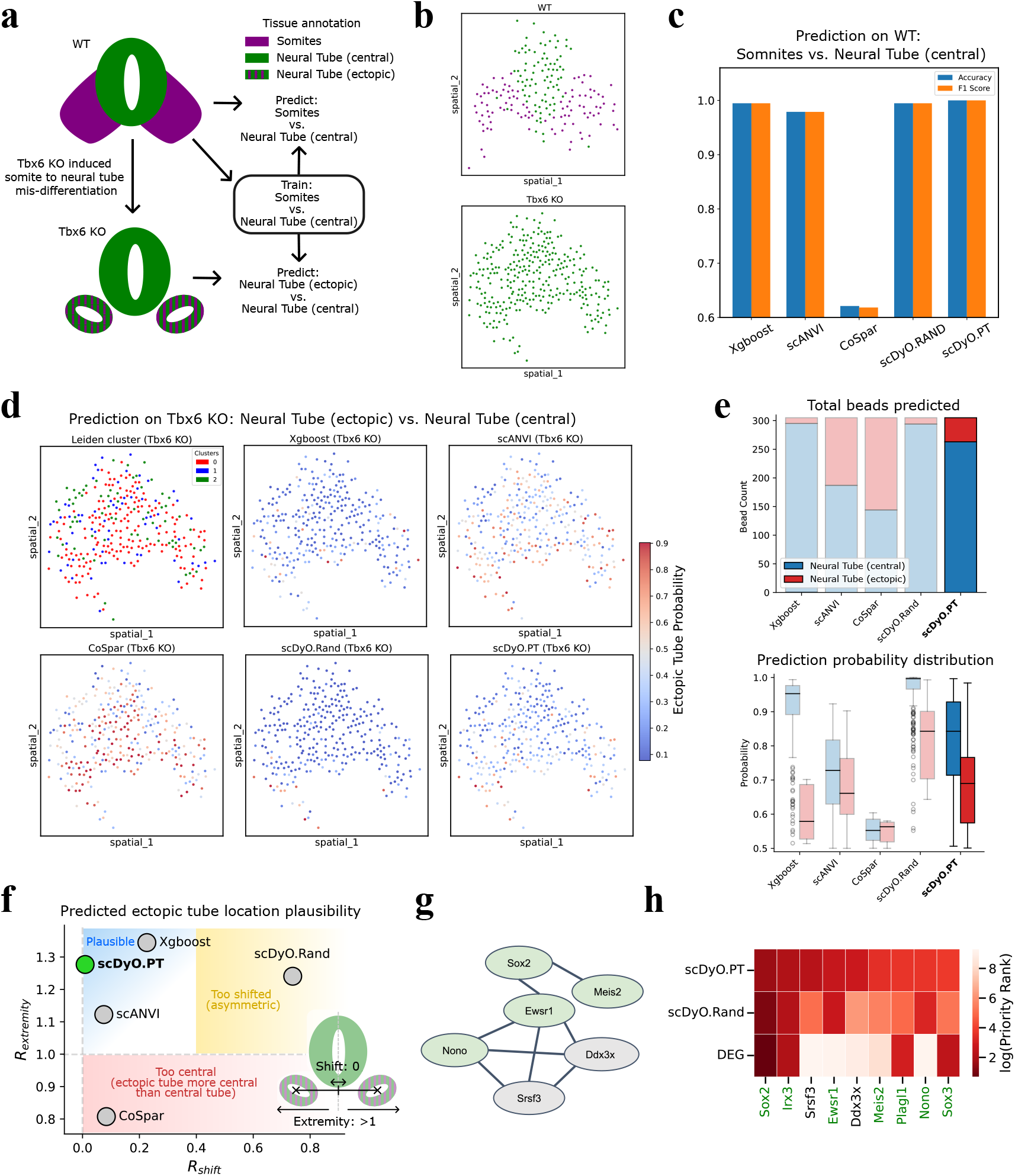
Reconstruction of spatial domains in genetically perturbed embryos. **a**, *Tbx6* knock-out (KO) leads to mis-differentiation of somite precursors to ectopic neural tube cells. After training on neural and somite expression profiles on a wild-type (WT) sample, models are tasked to classify expression profiles in a KO sample as neural or ectopic. **b**, Spatial distribution of Slide-seq beads and their assigned tissue annotations (color) for the WT and *Tbx6* KO slice. **c**, Classification metrics on the annotated WT slice for all benchmarked methods. **d**, Spatial distribution of predicted ectopic neural tube probability (color) in the *Tbx6* KO slice. scDynOmics recovers a more visible bilateral domain structure, whereas baselines produce sparse or spatially incoherent predictions. We also show a standard Leiden clustering result, which demonstrates the difficulty of defining coherent spatial patterns from the expression profiles. **e**, The number of beads predicted to be part of the central or ectopic neural tube (top) and the distribution of their prediciton probabilities (bottom) for all models. **f**, Quantitative evaluation of the predicted spatial distributions using the Extremity Ratio (*R*_*extremity*_) and Central Shift Ratio (*R*_*shift*_), see Methods. **g**, Key regulon for the ectopic neural tube domain reconstructed from the top 50 discriminative features identified by scDynOmics, utilizing the STRING[45] database. Documented transcription factors (TFs) are highlighted in green. **h**, Heatmap comparing priority rankings (color) of features (columns) between pretrained scDynOmics, non-pretrained scDynOmics, and differentially expressed genes (DEGs) from standard analysis (rows). While general neural markers (*Sox2, Irx3, Sox3*) are also found as standard DEGs, specific regulators like *Meis2* and *Ddx3x* are uniquely prioritized by the pretrained scDynOmics model.

To reconstruct the ectopic neural tube domains, we fine-tuned the scDynOmics model on the WT embryo slice containing annotated neural tube and somite populations, and subsequently applied it to the *Tbx6* KO slice. In the initial WT validation, pretrained scDynOmics achieved top-tier classification metrics, with XGBoost and the non-pretrained model trailing slightly behind (Fig. 6c). However, significant performance divergence emerged when reconstructing the ectopic domains in the KO slice. XGBoost and the non-pretrained scDynOmics model were overly conservative, annotating only a limited number of Slide-seq beads as ectopic neural tube and failing to form coherent spatial domains (Fig. 6d,e). Conversely, alternative methods like scANVI and CoSpar predicted a large number of beads with ectopic neural tube annotation but lacked clear spatial aggregation at the expected ectopic locations. In contrast, scDynOmics recovered spatially coherent populations of ectopic neural tube beads located bilaterally to the central neural tube, mirroring the expected spatial positioning of somites in a wild-type embryo (Fig. 6f).

Furthermore, we analyzed the key regulatory features driving scDynOmics to distinguish these ectopic domains from the central neural tube (Fig. 6g). While all of the pretrained scDy-nOmics, non-pretrained scDynOmics, and standard differential expression analysis consistently identified common neural markers like *Sox2, Irx3*, and *Sox3*, the pretrained model uniquely prioritized *Meis2* and *Ddx3x* (Fig. 6h). These findings are biologically significant: *Meis2* is well-documented as a critical regulator of central nervous system development[55, 56, 57], while *Ddx3x* is an essential regulator of neurogenesis[58, 59]. The ability of pretrained scDynOmics to prioritize these functionally critical factors, which can be missed by alternative analysis, demonstrates its capacity to extract important molecular signatures reflecting genetic perturbation.

## 3 Discussion

While scDynOmics effectively captures cellular states and regulatory logic, we identify few key avenues to extend its modeling capabilities toward more complex biological systems:

### 3.1 Scaling Pretraining with scRNA-seq Data

In the current manuscript we present scDynOmics only pretrained on paired scMultiomics data which constrains the pretraining corpus size due to the relative scarcity and high sparsity of such profiles compared to scRNA-seq. To leverage the vast size of pure transcriptomic atlases, future iterations will incorporate pretraining RNA velocity inspired multiomics corpora consisting of spliced and unspliced expression profiles derived from scRNA-seq datasets, similar to the fine-tuning application in Section 2.2. This strategy can significantly expand the scale of pretraining beyond limited multiomic resources.

### 3.2 Universal Cross-Species Tokenization

The current reliance on species-specific gene vocabularies restricts transfer learning across species. To engineer a more universal foundation model, one can adopt an ortholog-aware tokenization scheme, inspired by architectures like Nicheformer[60]. By learning a shared embedding space for homologous genes, scDynOmics could be pretrained on massive, multi-species atlases, enhancing generalization to data-scarce organisms by leveraging regulatory knowledge from multiple species.

### 3.3 Modeling Tissue-Level Interactions

The current framework treats cells as independent units, overlooking microenvironmental influences on cellular identity. Future work could extend the architecture to capture spatial dependencies via graph-based attention mechanisms or spatial positional encodings. This integration can enable the model to learn representations of tissue niches rather than isolated cells, unlocking new potential for dissecting organogenesis and tumor microenvironments.

### 3.4 Hardware-Aware Kernel Optimization

Although scDynOmics achieves algorithmic linear complexity via the Linformer mechanism and has a modest resource footprint (Supplementary Fig. 1), the computational efficiency could be further improved through hardware-aware optimizations. Future implementations could integrate optimized attention kernels, such as FlashAttention[27] or SAGEAttention[29], to minimize memory access overhead. Coupling our architectural optimizations with these kernel-level accelerations would allow for significantly larger batch sizes and faster convergence, maximizing the utility of available computational resources.

## 4 Conclusion

scDynOmics reconciles computational scalability with biological interpretability in single-cell representation learning. By embedding regulatory priors within a hybrid attention architecture, scDynOmics preserves the robustness and exceeds the performance of state-of-the-art baselines while decisively outperforming them in generating relevant insights in complex biological settings like developmental trajectories and perturbations. This unique capacity to uncover context-specific regulatory drivers, combined with coding-genome scalability and resource efficiency, establishes scDynOmics as a powerful framework for mechanistic discovery beyond the reach of existing approaches.

## 5 Methods

### 5.1 Linear Attention Formulation and Hybrid Encoders

Scaled self-attention in the standard Transformer model is formulated as:

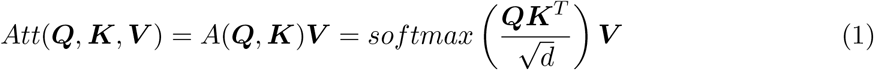

where ***Q, K, V*** ∈ ℝ^*L*×*d*^ represent the query, key, and value matrices for an input sequence of length *L* and embedding dimension *d*. Computing the full attention matrix *A* ∈ ℝ^*L*×*L*^ requires *O*(*L*^2^*d*) time and space, prohibiting its application to coding-genome scale inputs (*L* ≈ 20, 000).

To resolve this, scDynOmics integrates a Linformer-based low-rank approximation, projecting the *L* dimensional gene space to a lower-dimensional latent space *l*. Using two linear projection matrices ***E, F*** ∈ ℝ^*l*×*L*^, the Linformer approximated attention is defined as:

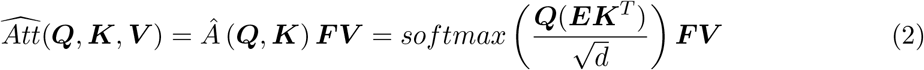

This factorization reduces the overall complexity to *O*(*lLd*). Taking constant *l* ≪ *L* and *d* ≪ *L*, Linformer can be considered having a linear complexity of *O*(*L*). This optimization can help to reduce memory and computational requirements for processing long sequences, significantly improving model scalability.

In other research domains, two major concerns have been raised with using Linformer: Linformer is not compatible with autoregressive processing that is essential for generative tasks in various domains, and Linformer assumes a fixed input length[61]. However, we posit these limitations are irrelevant for scDynOmics. Single-cell profiles lack a natural sequential ordering, and the coding-genome size *L* is fixed per species. Therefore, the fixed-length projection of Linformer provides a substantial efficiency gain without compromising the model’s ability to learn cellular semantics.

The scDynOmics encoder consists of a stack of modified Linformer blocks, each containing a multi-head self-attention (MHSA) module and a feed-forward network (FFN). To instantiate the biologically motivated architecture described in Section 2.1, we alternate between two distinct layer configurations: This hybrid approach further reduces computational cost relative to a fully unconstrained model while preserving robustness against incomplete biological annotations.

#### 5.1.1 TF-Encoders

In these constrained layers, the input sequence is explicitly subset to documented transcription factors (TFs) prior to projection such that ***E***, 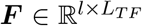, where *L*_*T F*_ represents the number of documented TFs[5]. This restricts the Key (***K***) and Value (***V***) generation to a subspace explicitly representing known regulatory drivers, forcing the attention mechanism to prioritize established biological networks and ground the embeddings in validated biology.

#### 5.1.2 Full-Encoders

In these unconstrained layers, the projection matrices ***E, F*** ∈ ℝ ^*l*×*L*^ span the entire coding-genome feature space. This enables the model to capture latent regulatory features and unannotated regulatory elements absent from the curated TF databases.

### 5.2 Input Embedding

Supporting the coding-genome scalability, the input cellular representations consist of a gene token sequence with fixed order according to the species and binned value token sequences based on the multimodal data. Denoting *v* as the multimodal data values, *m* as the modalities, and *t* as number of reserved tokens (e.g., `[MASK]`, `[PADDING]`), the embedding vector of gene *g* in a given input cell of species *s* can be obtained as:

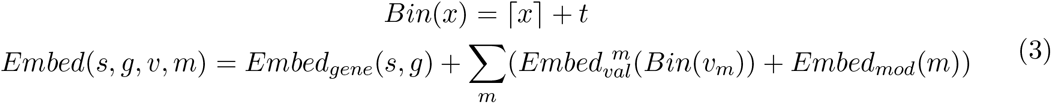

In the case of applying masking or padding on the input representation, associated tokens will be used to obtain corresponding embeddings.

### 5.3 Data Collection and Preprocessing

We assembled a raw collection of 5, 912, 810 mouse single-cell multiomic profiles from various public datasets, with a feature space comprising 22, 452 coding genes from the *Ensembl GRCm38* (mm10) assembly and documented TFs[5]. To ensure high-quality feature representation, RNA read counts were normalized using the transcripts per million (TPM) method and log-transformed. For the ATAC-seq modality, reads were assigned to genes if they mapped to *pan-promoter* regions, defined as the window extending from 1, 500 bp upstream to 500 bp downstream of transcription start sites (TSSs). Each gene’s mapped count was first divided by the number of associated TSSs, and subsequently normalized using the counts per million (CPM) method and log-transformed. Cellular samples were filtered following standard quality control recommendations[62], with the exception of lowering the minimum detected gene threshold to 100 to accommodate the inherent sparsity of scMultiomics data. As a result, 752, 155 single-cell multimodal profiles passing quality control in at least one modality were retained for pretraining (Fig. 2a).

For the experiments in Section 2.3, we additionally curated a distinct set of human sc-Multiomic PBMC and lymph node profiles with a feature space of 20, 492 genes based on the *Ensembl GRCh38* (hg38) genome assembly and documented TFs[5]. Adhering to the preprocessing and quality control protocols established for the mouse corpus, we retained a total of 48, 067 scMultiomic profiles.

### 5.4 Pretraining

Adopting the self-supervised learning paradigm of BERT[9], we employ a Masked Input Prediction (MIP) objective for pretraining (Fig. 1d). A key innovation in our approach is the structuring of input data to explicitly model biological dynamics. We configure the model to accept paired modalities, drawing a theoretical parallel to the concept of RNA velocity[6]. In velocity analysis, the ratio of unspliced precursor mRNA (pre-mRNA) to spliced mRNA provides a proxy for temporal changes in the RNA expression profile. In the scenario of scMultiomics data, the chromatin accessibility profiles of approximated promoter regions are analogous to the pre-mRNA profiles, while the gene expression profiles correspond to the spliced mRNA profiles. By training on these paired modalities, we encourage the model to learn the temporal and causal dependencies governing cellular state transitions.

During pretraining, we apply a masking strategy where a default ratio of 15% of genes with non-zero expression in at least one modality are randomly selected. Selected genes’ binned values are masked across all modalities. The model is then tasked with reconstructing these original values using the context provided by unmasked features. We employ separate linear reconstruction heads for each data modality to infer the original token bins from the encoder’s latent representations. The complexity of the reconstruction is deliberately constrained to compel the encoder to capture rich biological semantics, yielding expressive embeddings optimized for downstream adaptation. The reconstruction quality is optimized using a cross-entropy loss function calculated over the softmax probability distribution of the possible value bins.

### 5.5 Parameter-Efficient Fine-Tuning and Task Adaptation

To adapt the pretrained scDynOmics model to specific downstream tasks without incurring the computational cost of full-parameter updates, we employ a PEFT strategy. Instead of updating the entire network, we freeze the pretrained encoder and introduce lightweight, trainable adapter modules.

Let *W*_*n*_ ∈ ℝ ^*p*×*q*^ denote the frozen pretrained module of layer *n*, mapping input *x*_*n*_ ∈ ℝ ^*q*^ to a hidden vector *h*_*n*_ ∈ ℝ ^*p*^. Standard LoRA approximates the fine-tuning update Δ*W*_*n*_ with a low-rank decomposition Δ*W*_*n*_ ≈ *B*_*n*_*A*_*n*_, where *A*_*n*_ ∈ ℝ ^*r*×*q*^ and *B*_*n*_ ∈ ℝ ^*p*×*r*^ with *r* ≪ min(*p, q*). The layer output is therefore written as

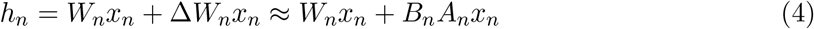

However, the standard LoRA is limited in approximating non-linear weight updates. In contrast, the adapter module[32] has been developed to introduce an additional non-linear activation function *Act* between the two low-rank projection matrices. With a skip connection over the low-rank update, the layer output is modified as:

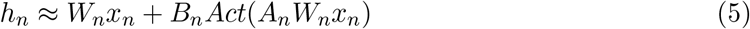

Instead of attaching to each linear weight, the adapter module is added only twice in a standard Transformer encoder layer: once after the MHSA block and once after the FFN block.

The architecture flexibly accepts monomodal or multimodal inputs, dynamically adjusting the relevant embedding components as defined in Eq. 3. The encoder weights associated with genes not involved in the downstream task, which do not appear in any modality of all cells in the query dataset, can be pruned for computational efficiency. The reconstructors in the pretraining process shall be replaced by task-specific modules, such as a classification model, according to the desired downstream task.

### 5.6 Gradient-based Interpretability Framework

To decode the biological logic driving scDynOmics’ predictions, we implement an attribution mechanism based on Integrated Gradients (IG)[63, 64]. Directly interpreting attention weights is not only unsuitable with scDynOmics’s PEFT strategy of tuning only adapter modules but also ignores the influences of downstream task-specific modules. In constrast, IG satisfies the axiom of completeness, ensuring that the attribution values sum to the difference between the model output and a baseline. We define the baseline reference as a zero-embedding vector, representing a theoretically “silent” cellular state with no read in any modality. Attributions are computed by backpropagating gradients through the full architecture as shown in Eq. 6. For a given input cell *x* predicted as class *c*, the attribution vector **IG**(*x, c*) quantifies the contribution of every feature to that specific prediction. To transition from local, single-cell level explanations to global regulatory signatures, we aggregate attributions across all *N* cells being predicted to a class *c*, yielding a mean attribution vector ***µ***_*c*_. To standardize these values across classes with varying prediction confidence, we apply *L*1 normalization, resulting in a relative importance score **s**_*c*_ that represents the proportional contribution of each input feature to the model output:

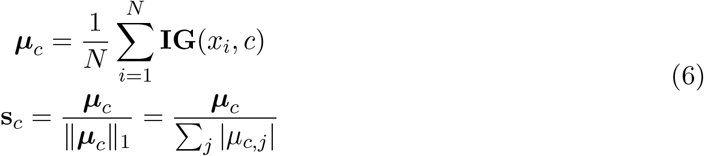

While **s**_*c*_ identifies important features, it does not distinguish between class-specific markers and ubiquitous “housekeeping” features that could be active across many cell classes. To isolate unique regulatory drivers, we introduce a discriminative specificity score (**d**_*c*_) especially for classification tasks. This metric penalizes broad signal by subtracting the maximum importance score observed for feature *j* in any background class *k* ≠ *c* from its importance in the target class:

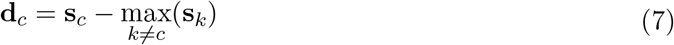

By filtering out broadly involved features, this formulation functions similar to an *in silico* differential analysis, amplifying genes that are uniquely predictive of a specific cell class while suppressing generic background signals.

### 5.7 Benchmarking against competing methods

All deep learning models, including scDynOmics, were trained and evaluated on a NVIDIA H100 80GB Tensor Core GPU leveraging CUDA acceleration. To ensure fair and rigorous comparison, hyperparameter tuning for baseline methods was conducted within the recommended ranges provided by their respective official guidelines. We maintain focus in the main figures by reporting the overall best-performing configuration across all experiments for each baseline.

#### 5.7.1 Comparisons against Single-Cell Foundation Models

For comparison against scBERT[17], we utilized the official scBERT repository and pretrained checkpoints provided by the authors. All data preprocessing and fine-tuning procedures were executed strictly using the default setting and hyperparameters recommended in the original implementation, ensuring a standardized baseline evaluation.

To benchmark against Geneformer[15], we retrieved the pretrained checkpoints for both version 1 (Geneformer.v1) and version 2 (Geneformer.v2.104M), the latter of which was pretrained on roughly 104 million transcriptomes. Fine-tuning was carried out following the authors’ guidelines in the Geneformer 0.1.0 release. Specifically, Geneformer.v1 utilized an input size of 2, 048 genes, while Geneformer.v2.104M utilized an input size of 4, 096 genes. Both versions were trained for a maximum of 20 epochs using the recommended default hyperparameters.

The CellFM[18] model was fine-tuned using the official CellFM-torch codebase and the published CellFM-80M checkpoint retrieved from the HuggingFace model hub. Because initial experiments revealed that the default mask ratio of 0.5 prevented the model from converging, the mask ratio was set to 0.0 to disable masking during fine-tuning. Furthermore, rather than using a fixed epoch limit, the model was trained with an early stopping criterion based on validation accuracy with a patience of 5 epochs. All other hyperparameters were kept at the default values recommended by the authors.

#### 5.7.2 Baseline Comparisons for Downstream Tasks

As a baseline for representative linear classification performance, we implemented a Logistic Regression model using the scikit-learn[65] library. To encourage feature sparsity and mitigate overfitting in the high-dimensional representation space, we applied an L1 regularization penalty. The model was optimized using the SAGA[66] solver, which is highly efficient for large scale datasets and compatible with L1 regularization. The inverse regularization strength (`C`) was set to 1.0, with a stopping tolerance of 1 × 10^−4^. The model was configured to calculate an intercept natively, utilizing a primal formulation and default intercept scaling, without applying predefined class weights.

To establish a robust, decision tree-based baseline, we implemented a gradient boosting classifier utilizing the XGBoost[38] framework. The model architecture was configured with a maximum tree depth of 6, and a learning rate (`eta`) of 0.1. To accommodate the multi-class nature of cell annotation and enable the calculation of threshold-independent evaluation metrics, the learning objective was fixed to `multi:softprob`. To ensure optimal convergence without arbitrarily limiting training duration, the maximum number of boosting rounds was expanded to 1, 000, governed by an early stopping criterion with a patience of 50 rounds.

scANVI[40], an established VAE-based model, was implemented utilizing the scvi-tools[67] library. The input `AnnData` object was configured via the `setup_anndata` function, explicitly registering the necessary batch covariates. The foundational scVI model was trained for a maximum of 300 epochs utilizing an early stopping criterion, after which the learned latent representations were extracted. Subsequently, the scANVI classification module was trained for a maximum of 100 epochs, also governed by early stopping to mitigate overfitting. By default, the latent space dimensionality was set to 30, with 4 hidden layers in total (`n_layers` = 2 for both the encoder and decoder). To assess the impact of structural complexity on classification performance, various scANVI configurations with differing hidden layer depths were rigorously evaluated. Specifically, the `n_layers` parameter was systematically varied from 1 to 5 for both the encoder and decoder. This resulted in models with total hidden layer counts ranging from 2 to 10, denoted as scANVI.L*n* where *n* ∈ {2, 4, 6, 8, 10}. Only the scANVI.L*n* models outperforming the default L4 configuration by the largest margin are shown in the main figures for clarity and focus on the most competitive baselines. Finally, feature explanation scores were extracted by enabling the `ig_interpretability` option during the scANVI prediction phase. The ranking of genes is obtained based on their associated attribution scores, which are calculated by the Integrated Gradients method[63].

#### 5.7.3 Cell Fate Prediction Benchmarks

For evaluating dynamic fate prediction capabilities, the CoSpar[52] framework was utilized to infer cellular transition maps as an OT-based benchmark. Temporal directionality was established by designating progenitor cells in Section 2.5 as the initial “early” time point, with the mature cells representing the “late” developmental stage. However, to apply CoSpar in predicting KO perturbation effects in Section 2.6, the KO condition was treated as the “early” state, while the control condition was treated as the “late” state. As explicit clonal barcoding was not utilized, the global transition map was inferred relying strictly on transcriptomic state information. This inference process was initialized utilizing an OT approach with Gene Expression Distance (GED) serving as the underlying cost metric. The algorithm was configured with a sparsity threshold of 0.2, employing the full similarity matrix alongside specific smoothing and iteration parameters. Following the construction of the transition map, progenitor fate biases were extracted for the designated terminal states. For downstream benchmark evaluation, deterministic lineage predictions were assigned to early-stage cells by selecting the terminal state with the maximum fate map transition score. Furthermore, normalized relative fate probabilities were computed for each cell utilizing the equation 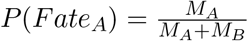, where *M* represents the raw transition map score for a respective terminal lineage.

As an additional robust baseline for trajectory inference and fate prediction, the CellRank[54] framework, integrated with the Moscot[68] toolkit, was utilized. Following standard preprocessing, the transcriptomic feature space was restricted to the top 2, 000 HVGs identified via the Seurat v3 algorithm implemented with Scanpy. Consistent with the CoSpar benchmark, temporal directionality was explicitly established by assigning progenitor cells to an initial time point and mature lineage cells to a subsequent time point. To model cellular dynamics over time, a Temporal Problem was formulated and solved utilizing Moscot’s Optimal Transport solver, configured with an entropic regularization parameter (*ϵ*) of 1 × 10^−3^ and an unbalanced transport penalty (*τ*_*a*_) of 0.95. The resulting transport maps were subsequently converted into a CellRank `RealTimeKernel`, and a transition matrix was computed incorporating all self-transitions and a connectivity weight of 0.2. To identify meta-stable lineage attractors, Generalized Perron Cluster Crossing Analysis (GPCCA) was employed, computing 20 Schur components to fit the macrostates. Terminal states were explicitly designated corresponding to the predefined mature somite and neural tube subpopulations. Finally, global fate probabilities were computed for each cell, and the absorption probabilities for the individual somite substates were aggregated to calculate a unified lineage probability. Deterministic cell fate predictions were assigned using a strict majority-probability threshold, whereby cells exhibiting an aggregated somite fate probability ≥ 0.5 were classified to the somite lineage, with the remainder assigned to the neural lineage as a standard binary classification task.

#### 5.7.4 Statistical Baselines for Key Feature Identification

In parallel with machine learning-based approaches differentially expressed genes (DEGs) were identified directly from the provided annotations to extract the defining gene signatures of each cellular subpopulation. These DEGs were computed utilizing the Wilcoxon rank-sum test via Scanpy’s `rank_genes_groups` module. Marker genes for each annotated category were identified and ranked based on their computed test statistic scores, rather than standard log-fold changes.

#### 5.7.5 Spatial Evaluation Metrics

To evaluate whether the predicted cell classes conform to the expected biological spatial priors of having neural tubes centrally distributed and ectopical tubes distributed bilaterally, we defined specific geometric ratios.

Let 𝒱 represent the set of all *N* = |𝒱| Slide-seq beads in the tissue section. The geometric center of the tissue along the spatial_1 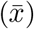 axis is inferred based on the mean spatial_1 coordinate of the beads located at the uppermost 5% quantile of the spatial_2 axis.

Let 𝒱_*c*_ ⊆ 𝒱 represent the subset of beads predicted as the targeted class *c*, where *n* = |𝒱_*c*_|, and *x*_*i*_ denote the spatial_1 coordinate of bead *i*. Conversely, let 𝒱_*others*_ = 𝒱 \ 𝒱_*c*_ represent the remaining beads in the tissue.

The **Extremity Ratio** (*R*_*extremity*_) measures the mean absolute distance of the predicted beads from the tissue center, relative to the remaining non-target beads:

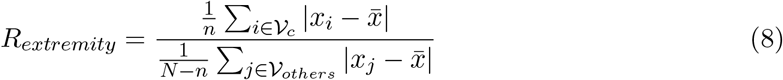

A ratio > 1.0 indicates the predicted beads are predominantly distributed at the spatial extremities of the tissue axis compared to the background tissue.

The **Central Shift Ratio** (*R*_*shift*_) quantifies the absolute deviation of the predicted beads’ mean spatial coordinate from the tissue center, normalized by the location of the center coordinate:

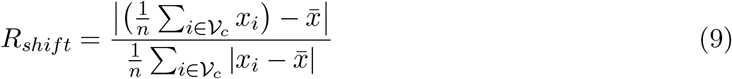

This ratio evaluates the bilateral symmetry of the prediction; values deviating further from 0 indicate that the target class population is asymmetrically shifted to one side, left or right, of the central axis.

### 5.8 Implementation Details

The scDynOmics framework was implemented with Pytorch Lightning[69]. To initialize the model parameters, the weights of all linear layers were drawn from a normal distribution with a mean of 0.0 and a standard deviation of 0.02, while their corresponding bias terms were strictly initialized to 0.0. For all layer normalization components, the scale (weight) and shift (bias) parameters were initialized to 1.0 and 0.0, respectively. Embedding layers were purposefully excluded from this custom scheme, relying on standard PyTorch[70] initializations. The default precision setting for all computations was 32-bit floating point (FP32). For the selected L12H12 architecture, the input embedding dimension was set to 96 while the FFN dimension was set to 768.

All pretrainings are performed on a single compute node with 8 NVIDIA H100 80GB Tensor Core GPUs, with a default 32-bit precision settting. We use the AdamW optimizer[71] with `betas` 0.9 and 0.98, weight decay of 0.01 and dropout of 0.0. The batch size is set to 240 with each device processing 6 samples. The learning rate was modulated using a cosine annealing schedule with warm restarts[72, 70], decaying from an initial value of 5 × 10^−5^ to a minimum of 5 × 10^−6^ over a primary cycle of 20 epochs, with the duration of each subsequent cycle doubling. Checkpoints were taken every epoch. The mouse model was pretrained for 10 epochs (~30,000 steps) for ~24 hours, while the human immune cell model was pretrained for 100 epochs (~20,000 steps) for ~15 hours.

For all fine-tuning experiments, the batch size was uniformly maintained at 16. We retained the AdamW optimizer configuration utilized during pretraining. The learning rate strategy featured a one-epoch warmup, initiating at 2 × 10^−4^ and peaking at 2 × 10^−3^. Subsequent to this warmup, a modified cosine annealing schedule was applied, decaying to a minimum learning rate of 1 × 10^−4^ over a constant cycle length of 5 epochs. To mitigate overfitting, early stopping was implemented with a patience of 5 epochs based on validation loss. For LoRA adapter modules, the activation function was set to ReLU. Prior to downstream classification, max pooling across the embedding dimension was applied to the encoder outputs. By default, the LoRA adapter latent dimension was set to 4, coupled with a downstream multi-layer perceptron (MLP) with a single hidden layer and an output layer, featuring a hidden dimension of 256 and ReLU activations. However, for larger, more complex fine-tuning datasets (i.e., Section 2.2, Section 2.3), the adapter dimension was expanded to 8, and the downstream MLP hidden dimension was increased to 640.

Aside from benchmarking the computational resources for fine-tuning on the NVIDIA H100 (Supplementary Fig. 1), we also evaluated usability with limited computational resources. For this we also performed the same benchmarked fine-tuning task on a single consumer grade NVIDIA RTX 5090 32GB GPU with a batch size of 32, finishing fine-tuning in 74 ± 5 minutes (mean ± standard deviation over 10-fold cross validation).

To systematically evaluate the performance gains conferred by pretraining, we compared the pretrained scDynOmics model against a non-pretrained baseline (scDyO.RAND). Both model variants were evaluated using identical architectures and fine-tuning configurations, differing exclusively in whether after random initialization their encoder weights were pretrained or not.

### 5.9 Data Presentation

All box plots show the interquartile range as the box, the median as line in the box, the full range of the distribution except outliers as whiskers, and the outliers defined as all datapoints outside of 1.5 times the interquartile range from the interquartile range as circles.

## 6 Data availability

The raw mouse scMultiomic datasets curated for this study were obtained from various public sources. These data can be accessed via the Gene Expression Omnibus (GEO) under the following accession numbers: GSE253345[73], GSE180900[74], GSE205117[75], GSE231674[76], GSE184981[77], GSE192788[78], GSE211464[79], GSE210234[80], GSE209610[81], GSE126074[82] (all 34 mouse samples), GSE211542 paired with GSE211543[83], GSE140203[4] with 4 mouse samples (GSM4156597 paired with GSM4156608; GSM4156599 paired with GSM4156610), GSE117089[84] (GSM3271044 paired with GSM3271045), GSE130399[85] (4 mouse samples: GSM3737490; GSM3737495; GSM3737496; GSM3737499), GSE229513[44] with 2 mouse samples of GSM7164730 and GSM7164731 while 1, 000 cells in each sample were randomly held out from the pretraining process for the model explanation experiment in Section 2.4. The E-MTAB-11264[86] dataset can be accessed through the ArrayExpress database. Furthermore, the following datasets can be accessed via 10x Genomics Datasets portal: Alzheimer’s Disease Mouse Model Brain, Mouse Kidney Nuclei, Mouse Brain Nuclei, and Fresh Embryonic E18 Mouse Brain. The scRNA-seq mouse gastrulation dataset[37] is processed into a multimodal RNA velocity dataset with the scVelo[87] pipeline and directly available through the scVelo package. All of the spatially resolved Slide-seq datasets in studying mouse embryo development and *Tbx6* KO effects[51] are available through the CELLxGENE database.

The raw human scMultiomic immune cell datasets were obtained from the 10x Genomics Datasets portal: Lymph Node with B Cell Lymphoma, Human PBMCs via Chromium X, Human PBMCs via Chromium Controller, Healthy Donor PBMCs (3k), Healthy Donor PBMCs (10k). The annotated and preprocessed Zheng68k[41] dataset is available via GitHub[88].

## 7 Code availability

The source of scDynOmics is available on Github (github.com/KlughammerLab/scDynOmics), the Python package on PyPI, and the documentation on GitHub Pages including tutorial notebooks for downstream applications as well as instructions for extending pretraining to new datasets.

## 8 Author contributions

G.Y. conceived the project with contributions from J.K. and S.W.M.. G.Y. implemented models, curated data, and analyzed data. G.Y. designed visualizations with contributions from T.J.S.R, J.K. and S.W.M.. J.K. and S.W.M. supervised the research. G.Y., J.K. and S.W.M. wrote the manuscript with contributions from T.J.S.R.. All authors read, corrected, and approved the manuscript.

## 9 Acknowledgements

The authors thank Jan Watter for assistance with publishing the package on PyPI, and Karl-Peter Hopfner and Gregor Witte for providing and maintaining local GPU compute resources. The authors gratefully acknowledge LMU Klinikum for providing computing resources on their Clinical Open Research Engine (CORE). The work was supported by German Research Foundation (DFG) grants CRC237 369799452 (J.K.) and CRC274 408885537 (J.K.), an Else-Kröner-Fresenius-Stiftung starting grant 2019_A70 (J.K.), and by LMUexcellent (S.W.M.), funded by the Federal Ministry of Education and Research (BMBF) and the Free State of Bavaria under the Excellence Strategy of the Federal Government and the Länder.

## Supplementary

**Supplementary Fig. 1:**
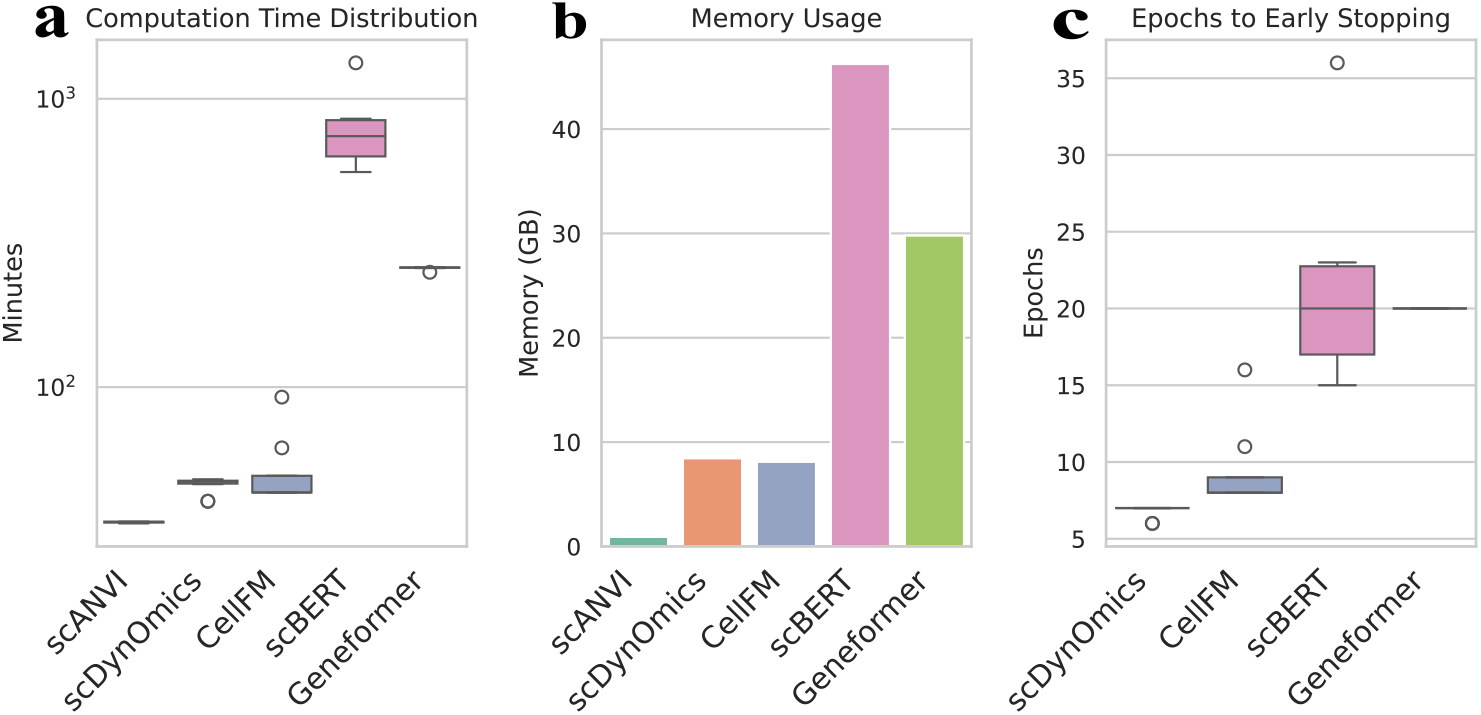
Computational performance benchmark with the Zheng68k dataset. **a**, Ten-fold cross-validation runtimes on one NVIDIA H100 GPU comparing scDynOmics to the baseline scANVI and three existing foundation models (CellFM, scBERT, Geneformer). All models except scANVI were run with a per-device batch size of 8. All models except scANVI and Geneformer were fine-tuned with an early stopping patience of 5 epochs; Geneformer was trained for a fixed 20 epochs. Note that by scaling the per-device batch size by a factor 2^*n*^, the memory of scDynOmics scales roughly the same while the runtime scales inversely, enabling its flexible optimization to diverse computing setups. **b**, Peak GPU memory usage during training for all models. **c**, Number of epochs required for fine-tuning of scDynOmics and other foundation models.

